# Evolutionary macroecology: Incorporating phylogenetic information more explicitly into process-based, forward time, simulation models

**DOI:** 10.1101/2024.10.18.619050

**Authors:** Victoria Culshaw, Thiago F.L.V.B. Rangel, Isabel Sanmartín

## Abstract

A conceptual and mechanistic approach for bridging the fields of macroecology (ecological biogeography) and historical biogeography has been a long-term aim in Evolutionary Biology. Such a bridge could increase our understanding on the processes governing the spatial and temporal generation of biodiversity distribution patterns. This aim has been recently approached from the perspective of evolutionary biogeographic inferential statistical models, within a maximum likelihood or Bayesian framework, which incorporates the contribution of environmental factors as scaling parameters. Here, we describe a new approach that builds on a spatially-explicit, forward-time, computer simulation (“automat”) model. The model sets a series of rules by which speciation, extinction, and dispersal of lineages are governed within an environmentally heterogeneous, two-dimensional gridded landscape. Unlike some previous approaches, niche conservatism is assumed but the model allows for environmental conditions to vary both spatially and temporally, by letting the model run over three time-series of actual palaeoclimate data, spanning the last 20 million years of geological history. Speciation is governed by a global speciation rate, whereas the background extinction rate is made dependent on abiotic (palaeoenvironmental conditions) and biotic (species density per grid cell) factors, hence giving a local background extinction rate. Also, we propose a novel mechanistic approach in which species are not the result of unique, independent events but linked through evolutionary history from a single evolutionary origin. We set different rules to generate the resulting phylogenies to test different factors (time, environment) that govern the inheritance of range distributions. Dispersal follows a simple Poisson kernel model, with higher probability of migration to contiguous grid cells and rare long-distance movements to distant cells. We also describe ways in which the presence of temporal dispersal barriers could modify the resultant spatial patterns. Evaluation of model accuracy and fit is based on comparison of simulated spatial patterns with observed, empirical ones. We use statistic dependent variables such as the spatial distribution of species in the landscape, species’ geographic range size and location, and the shape of the resultant phylogenies. Finally, we propose that this spatially-explicit simulation model could be used to evaluate the role played by niche conservatism, ecological vicariance and climatic-driven extinction in the generation of disjunct continental patterns, such as the presence of sister-lineages in numerous families of angiosperms forming a ring in the continental margins of the African continent, the Rand Flora pattern.

## Introduction

Understanding biodiversity patterns –why some biotas are more or less diverse than others and how local or regional biotas became assembled– has become a pressing goal in biodiversity research in light of the Anthropocene biodiversity crisis (up to 25% of extant species will become extinct in the next decades; IPBES, https://ipbes.net) and ongoing global climate change scenario (linked to human activities). Global biodiversity patterns, such as the Latitudinal Diversity Gradient (LDG) –the observed general increase in species richness or biodiversity from the poles to the tropics (Fischer 1960; Pianka 1966; Stevens 1989)– have traditionally been explained in relation to shifts in environmental variables such as temperature or precipitation (Hillebrand 2004; Kreft and Jetz 2007). There is, though, a current understanding that environmental variables cannot by themselves increase or decrease local or regional species richness; evolutionary processes such as dispersal, extinction, or speciation must be taken into account (Qian and Ricklefs 2004; Wiens and Donoghue 2004; Jablonski et al. 2008; Mittelback et al. 2007). Thus, understanding the mechanistic, causal factors driving speciation, dispersal, and extinction rates in relation to fluctuating spatially and temporal environmental (climatic) conditions should allow us to comprehend why some biotas are richer or more species-poor than others or how global biodiversity patterns have been shaped over time. For example, it has been hypothesised that if niche conservatism prevails over niche evolution in regions of spatially heterogeneous and temporally fluctuating climate, diversification will occur predominantly by a process of range fragmentation, caused by the inability of species to adapt to changing environmental conditions in portions of the ancestral range, in what has been termed the process of “ecological vicariance” (Wiens 2004; Wiens and Donoghue 2004; Rangel et al. 2007).

These questions have been typically approached from two different perspectives. One of them is macroecological, by linking measures of diversity and spatial patterns of species richness to environmental (often, climatic) variables through “curve-fitting” correlative approaches, such as generalised linear modelling techniques (GLM; Hawkins et al. 2003; Rahbek et al. 2007; Chefaoui and Lobo 2007; Mairal et al. 2017), or through more complex spatial Ecological Niche Models (ENM) and Species Distribution Models (SDM, see Owens et al. 2011 for distinction between ENM and species distribution models, SDMs, from here we consider ENM only; Thuiller 2003; Araujo et al. 2005; Thuiller et al. 2006; Lobo et al. 2008; Hengl et al. 2009; Culshaw et al. in 2021b.). ENMs are a group of distances or models that are used to describe and predict climatic habitat suitability species’ geographical ranges from occurrence, and if possible, absence data, e.g. ecological niche factor analysis (ENFA; Hirzel et al. 2002; Basille et al. 2008; Hengl et al. 2009) or Mahalanobis distances (MD, Clark et al. 1998). ENMs for individual species can be overlaid to extract general patterns on climatic habitat suitability (Culshaw et al. in 2021b) These models have later been extended with the addition of other parameters, such as species interactions, non-climatic predictors, or spatial autocorrelation terms (Araújo and Luoto 2007; Dormann et al. 2007; De Marco et al. 2008; Guisan and Rahbek 2011). Currie et al. (1999) criticised curve-fitting approaches because they often do not provide quantitative predictions and because they “implicitly” assume that species richness patterns depend only on environmental variables, ignoring the contribution of historical and evolutionary events to such patterns. These critiques can also be extended to ENMs: overlapping of potential or realised species ranges in a gridded map does not provide a causal explanation behind the observed pattern.

The second approach to tackle the origin and maintenance of spatial patterns of diversity is using methods stemming from the fields of phylogenetics and historical biogeography (Sanmartín 2014). These methods use phylogenetic data (evolutionary relationships) and divergence time estimates, together with species occurrence data, to infer the distribution range of a species’ ancestors and the rate of evolutionary processes like speciation, extinction, and dispersal (Wiley 1981; Brooks and McLennan 1991; Ronquist and Sanmartín 2011; Sanmartín 2014). Current biogeographical inferential approaches are based on probabilistic models of range evolution (continuous-time discrete-state Markov chain models) and use the statistical maximum likelihood framework or Bayesian Inference to estimate ancestral states and parameter values. Unlike in ENMs, areas in biogeography are often defined in abstract terms, without an explicit spatial or environmental component. However, as some biogeographers have noted (Brooks 1990; Cracraft 1994; Sanmartín et al. 2001; Wiens and Donoghue 2004; Meseguer et al. 2015), the environmental (climatic) preferences of a species plays a strong role in constraining evolutionary and biogeographical processes such as the rate of ecological speciation, local background extinction and geographic expansion, and therefore should be considered in the inference of spatial patterns of species richness.

Understanding the mechanistic basis of broad-scale biodiversity patterns formed by persistence, adaptation, extinction or geographical range shifts remains the holy grail of modern biogeography and macroecology (Judson 1994; Willig et al. 2003; Gotelli et al. 2009). In the early days of these disciplines (XIX century and early XX century), phylogeny and biogeography were closely interlinked in the works of taxonomists and naturalists (Cox, Moore and Ladle. Biogeography: An Ecological and Evolutionary Approach, 10th edition, 2016). However, for over the last 40 years, the field of biogeography has been mainly concerned with building phylogenies and inferring the spatial-temporal evolution of lineages, with ecology deemed moot (Wiley 1981; Brooks and McLennan 1991; Ree and Smith 2008; Sanmartín et al. 2008; Lemey et al. 2009); conversely, macroecology has been slow in integrating the evolutionary perspective (Hawkins et al. 2003). Yet, historical biogeography and ecology have much to learn from one another (Wiens and Donoghue (2004); Hawkins et al. (2005); Culshaw et al. (2021a and 2021b), and an approach that bridges these two fields into a single theoretical and analytical framework will be crucial in exploring the relationship between ecological and evolutionary processes in the shaping of Earth’s species geographic ranges (Ricklefs 2006).

Recently, there have been some attempts to create this bridge. Studies such as Evans et al. (2009), Richards et al. (2007) or Meseguer et al. (2015) have championed the integration of phylogenetic and biogeographical inference with ENM frameworks. Richards et al. (2007) used hindcasted ENMs to inform coalescent simulations and test alternative coalescent hypotheses on population spatial dynamics. Evans et al. (2009) employed ENMs to quantify the climatic disparity among taxa and obtain an understanding of the degree along which niches evolved, and used this information as a secondary calibration to predict niche occupancy through the timeline of a dated phylogenetic tree. Meseguer et al. (2015) combined fossil record information with presence only data to infer the climatic tolerances of ancestral lineages and generate hindcasted ENMs. Subsequently, they used these models, as well as the spatial information provided by fossils, to inform probabilistic biogeographic inference about the existence of temporal climatically suitable corridors and climatic dispersal barriers. We recently proposed an approach (Culshaw et al. 2021b) to combine the biogeographic and ENM frameworks as complementary, independent sources of evidence, allowing the user to integrate the geographical information without sacrificing the evolutionary information, and vice versa. Hindcasted ENMs were used to inform biogeographic inference along long branches lacking cladogenetic events, while inferred ancestral ranges were used to select the truncation threshold needed to convert continuous habitat suitability values returned by the ENM into presence-absence predictions.

Though these integrative/combined approaches can improve the accuracy of the inferred biogeographic scenarios or the parameter estimates (Meseguer et al. 2015; Culshaw et al. 2021b), they are more geared towards understanding the spatio-temporal history of particular lineages and less to the testing of hypotheses on macroecological patterns. Biogeographic and ENM approaches do not allow us to examine the causes behind an observed correlation between species distribution patterns and environmental factors, or what this relationship would be like under small variations in the primary data. Knowing how stable a statistical relationship is, provides a better understanding of its predictive power in forecasting future scenarios. A mechanistic understanding of species distribution patterns requires modelling the processes by which these patterns are formed (Gotelli et al. 2009).

A third avenue to combine biogeography and macroecology is based on the use of spatially explicit, forward-time computational general simulation models (GSM, Rangel et al. 2006, 2007; Rabehk et al. 2007; Colwell et al. 2009; Gotelli et al. 2009), sometimes coined as “automat” or “in silico” models. In its simplest form, GSMs are mechanistic models that are governed by a set of rules for location; probability and mechanism of speciation; the inheritance of niche characteristics by each new species from its immediate ancestor; and the species ability to disperse to and successfully colonise new grid cells based on their environmental characteristics (Rangel et al. 2007; Gotelli et al. 2009). These sets of rules describe how a species may speciate, disperse and become extinct in an environmentally heterogeneous landscape (represented as a gridded domain; Gotelli et al. 2009). Typically, these models are probabilistic or stochastic (an example exception would be Hassell et al. 1991), and are run for a given number of time steps *t* or until a specific condition is met (e.g. until a particular number of species ranges are simulated or until a balance between speciation and extinction is achieved) and can accommodate for contemporary, past or future climates, evolutionary and historical forces, and geometric constraints. These models build frameworks to investigate hypotheses about the relative influence on species richness played by geometric dispersal constraints, environmental factors and historical processes (Rabhek et al. 2007; Gotelli et al. 2009), making these models more than suitable for exploring the stability of an empirical relationship. See Gotelli et al. (2009) for an excellent review on macroecology simulation models.

### Spatially explicit general simulation models (GSM)

Exploring the causes of spatial variation in species distributions through computational general simulation models (GSM) has been the subject of a rich literature. The GSM relies only on two data layers: a gridded map of a region or biogeographic domain with cells of a certain resolution; and the environmental variables measured on each grid cell. Unlike ENMs, GSMs do not require species occurrence data. Most GSMs include also three components or control knobs for simulating species richness patterns on the gridded domain: dispersal limitation, environmental gradients, and evolutionary events (Gotelli et al. 2009).

*Dispersal Limitation:* Levins (1969) was one of the first models that explored the introduction of spatial parameters into population models. Levins’s model describes a gridded landscape in which present in each cell is an unstable population system. Here a population becomes extinct within a specified number of time steps. However, when several populations are linked together through a spatial parameter modelling dispersal, the populations only face temporarily extinction as they can be re-colonised by another population, hence forming a stable “on our own we are weak, together we are strong” system, provided that the population models’ extinctions are in a state of asynchrony to one another. Levins’s model was very simple, assuming the landscape to be homogenous, and dispersal to be limitless to any cell within the gridded domain. Also the model only recorded whether a cell was occupied or not, i.e. number of individuals of the population within the cell was disregarded. This model of linked populations became known as the Levins metapopulation model, and it is widely regarded as the null model in exploring spatial patterns through GSMs (Gotelli et al. 2009). Further advances from these first models have included the “spreading dye model”, which imposed limitations on how a species may disperse through the landscape from its original cell: i.e. dispersal is determined by distance to the adjacent grid cells (e.g. Hassell et al. 1991; Jetz and Rahbek 2001; Connolly 2005).

*Environmental gradients* are typically based on temperature and precipitation measurements for each cell in the domain, and usually represent present-day climate data, and, increasingly popularly, palaeoclimate reconstructions (Meseguer et al. 2015; Culshaw et al. 2021b). With the use of ENMs, these environmental gradients can be converted into species-specific, habitat suitability values, where one of these suitability values is allocated per cell in the domain. Converting environmental variables into habitat suitability variables can be advantageous. One advantage would be that habitat suitability values from the ENMs can locate the potential spatial barriers to the spatial distribution of a population through time, and the GSM can be used to see whether it is possible for the population to overcome these spatial barriers given enough time, hence effectively mechanising the ENM (Thuiller et al. 2008). A second advantage is that speciation, background extinction and colonisation success rates can be converted by the ENM habitat suitability values from being global rates that are equal across species to local and species-specific.

Including *evolutionary events* in GSMs has mainly been tackled in one of two ways: i. by specifying a priori the number of independent evolutionary events that should take place during the simulation (e.g. Grytnes 2003; Storch et al. 2006; Rahbek et al. 2007); and ii. by including some scale or logic measurement of what a species is, running the simulation for a given time, and then observing how many evolutionary events took place and how many species are present at the end of the simulation (e.g. Rangel et al 2007; Boone 2010). In the first case, each species has an independent evolutionary origin and, if the niche is considered, the niche of each species is independent from the niche of other species; in the second, all species are linked by a single evolutionary origin, an ancestral species, and the sequence of speciation events depends on the set of rules that determines the balance between niche conservation and niche evolution (Rangel et al. 2007).

Evaluation of the power of GSMs is based on comparison of simulated and observed patterns, and the ability to predict or recover system-level properties such as the LDG pattern or the observed mid-domain effect (Gotelli et al. 2009). Comparison between simulations and the observed input data can also be used to find the optimal parameter values: these are the parameter combinations that maximise the similarity between the observed and the simulated patterns, for example, in regards to species richness and/or the geographic range distribution frequency among species (Rangel et al. 2007). In fact, one advantage of GSM approaches, which resembles that of Approximate Bayesian Computation methods in evolutionary biology (Leuenberger and Wegman 2010), is their flexibility to incorporate new parameters or variables to increase the realism of the model, since it becomes unnecessary to define the mathematical functions governing parameter behaviour/relationships analytically (i.e., likelihood-free methods).

### Some Limitations of GSMs

One of the main limitations of GSMs is the assumption of species equivalency. As in MacArthur and Wilson’s (1967) Island Biogeography Theory, or Hubbell’s (2001) Neutral Theory, rates of speciation, background extinction, and dispersal do not depend on species-specific characteristics, but are assumed similar or equal across species. Model realism could also be increased by incorporating actual palaeoclimate data and a real time scale in the simulations, rather than using an artificially fluctuating environment (Rangel et al. 2007). To consider speciation events as independent evolutionary events follows the classic ecological approach of using numbers of species or species richness to detect patterns (Jetz and Rabnek 2001; Wiens and Donoghue 2004), but does not help understand how these numbers are generated (Wiens and Donoghue 2004). Modelling evolutionary events using a scale and logic measurement of what a species is through a set of rules (Rangel et al. 2007) allows comparing the observed species richness patterns and geographic range size variation with the one obtained in the simulations, and hence to extract conclusions on the underlying processes. Yet, deciding upon this scale and logic measurement can be difficult, as the initial empirical ecological niche, the empirical heritability of the ecological niche from ancestor to descendant taxon, and an understanding of the balance between empirical niche conservation and empirical niche evolution through time must be known. This difficulty increases the appeal of removing oneself from the real world (such as done in studies like Rangel et al. 2007), and hence remove the need for empirical information and allowing the luxury of choice as to which type of rules are set. However, with this entrance into the virtual world comes a new challenge, the lack of an evolutionary component provided by a real phylogeny, for example, the inheritance of the ancestral niche by the descendants. Setting the simulation so as to explore the relationship between a particular phylogeny, or empirical species relationship, with a distribution pattern, climate history and niche dynamics allows testing whether a similar relationship could be recreated under different environmental conditions and niche dynamics. In other words, are the relationships we find within a phylogenetic tree linked to the distribution patterning or could the same relationships have been achieved through different distribution patterns, and vice versa?

Here, we describe a novel general simulation model which proposes a different way of integrating evolutionary history in GSMs in the landscape. The model includes a dispersal kernel to navigate around temporary barriers, and information on the tree and speciation intervals among species, so that the evolutionary (speciation) events in the simulation are dependent on one another, i.e., they have a single origin. We gain the information for this new evolutionary component by including phylogenetic information into the simulation. Unlike in Rangel et al. (2007), niche conservation is assumed within our model (i.e., no information on niche evolution); hence, evolutionary events are initiated by a speciation rate rather than by the balance between niche conservatism and evolution in a fluctuating environment. Moreover, our GSM uses time series of paleoclimatic data: in particular, three paleoclimate layers representing major cooling and warming events in the last 20 million years. As in other GSMs, the model uses a pattern-oriented modelling approach to compare the fit between observed and simulated data.

## A Novel General Simulation Model Introducing Evolutionary History Model Overview

Our new model follows Wiens and Donoghue (2004) biogeographic framework, in which the geographical range of a clade is determined by i. the ancestral ecological niche of the clade; ii. the geographical dispersal origin; iii. dispersal limitation described by abiotic and biotic conditions (e.g. habitat suitability and competition); iv. the potential for niche evolution presented by the geographical range, if niche conservation is not considered), and the length of time elapsed since the origin of the clade for niche evolution (if niche conservation is not considered) and dispersal to occur. Our spatially explicit, forward time GSM model directly addresses four of these points (i-iii and v). Point iv (the potential for niche evolution) is not addressed; since we assume niche conservatism or at least that we lack enough information to infer niche evolution. The model is built upon an n-dimensional niche space and a two-dimensional geographical, gridded cell map space. Each cell contains information on the number of species present within it, but not the number of individuals for each species, as well as its geographical coordinates, and by the local values of the same n environmental parameters that define the niche space (a representation pioneered by Pulliam 2000). The model has been written in the programming language PASCAL, and is available for distribution by demand to main author.

Figure 1 shows a scheme representing the model, showing the three components or control knobs in the model (Figure 1 the vertices of the triangle), and the relationships among these components in the model (Figure 1 the sides of the triangle). The three control knobs are the: a. Evolutionary Origins: including the rates of extinction (*μ*) and speciation (*λ*), and the time intervals between speciation events and/or the species tree topology, which provides information as to when a speciation event takes place and to whom this speciation event is related. b. Suitability: provides information on the abiotic and biotic suitability of each cell. Abiotic suitability defines how suitable the cell is for the species, given its climatic niche breadth and the measurements of abiotic variables, such as mean temperature and precipitation, in the cell. Biotic suitability refers to competition within each cell and is represented simply by the number of species present within a cell (i.e., density-dependence effects). c. Range Expansion: describes the ability of a species to disperse across the landscape. We used a model that combines the “spreading dye model”, allowing local dispersal to adjacent cells within a maximum of distance *M*, with rare events of long-distance (“freak”) dispersal to cells located beyond M.

**Figure 1.**
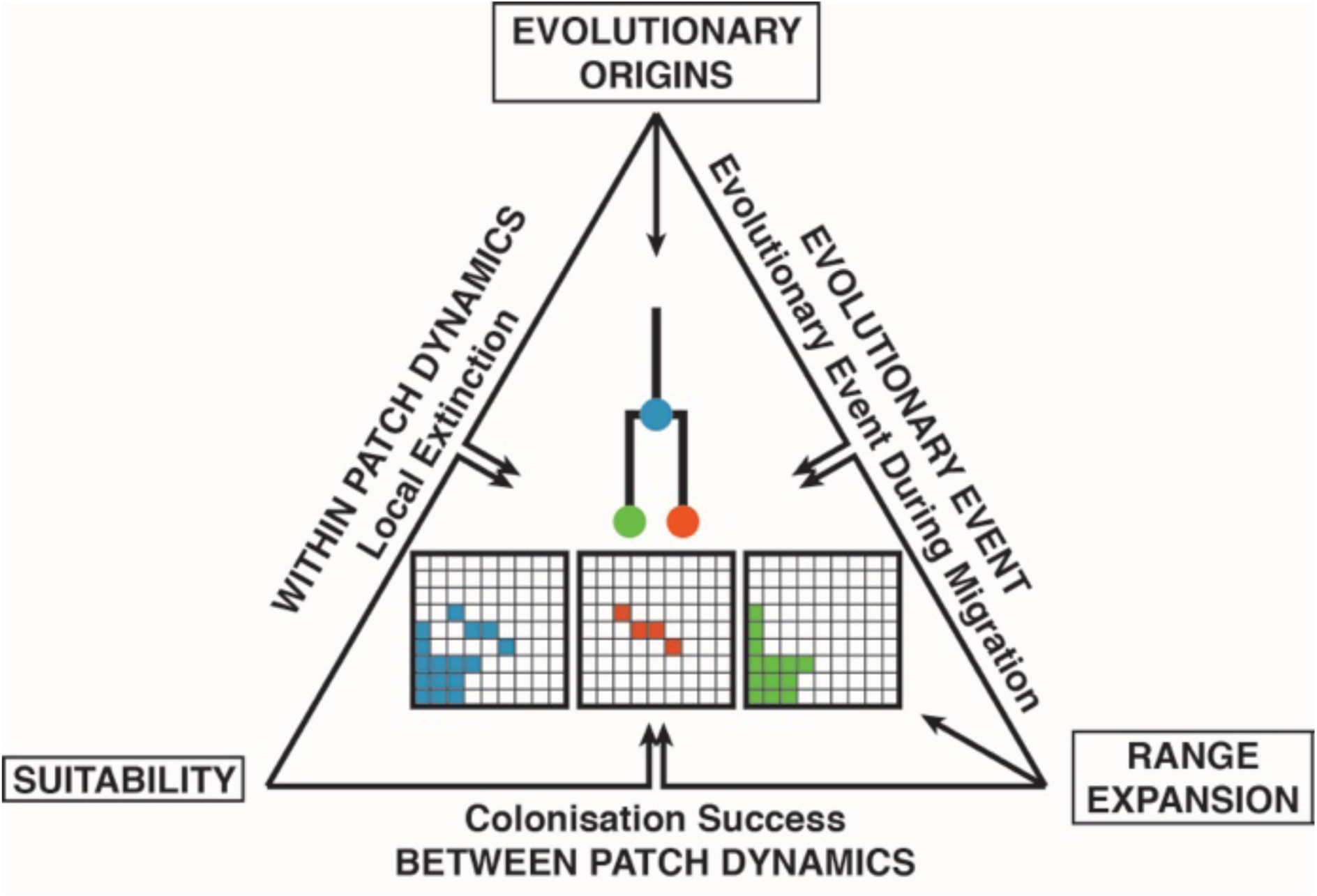
Model Overview. The points of the triangle represent the data layers supplied to the model. The triangle sides represent the connecting relationships between these layers. The black arrows represent the direct influence over the centre model results. Notice that “Suitability” only affects the model indirectly through “Within Patch Dynamics” and “Between Patch Dynamics”.

Relationships between components in the model (Figure 1, represented by the triangle sides) are: i. Within Cell Dynamics: models the effect of cell suitability on the local extinction rate to calculate the probability of species prevalence at time *t* within a cell. ii. Between Cell Dynamics models the colonisation success to a cell of a dispersing species through the relationship between cell suitability and dispersal success. iii. Evolutionary Events: models how the speciating species’ range is divided up between its daughter lineages through the relationship between the speciation rate, the dispersal kernel, and the environmental characteristics of the cells (i.e. habitat suitability). Niche conservation, and constancy of speciation and background extinction rates are assumed to occur between and within species, but not necessarily between time steps.

## Model Algorithm

In this section, we describe in more detail the model algorithm specifying the set of rules and functions that link the parameters in the model, as well as the distribution probability of these. Figure 2 shows an example of how the model algorithm works for one-time step *t* in which there is one event of long distance dispersal and one successful evolutionary event.

**Figure 2.**
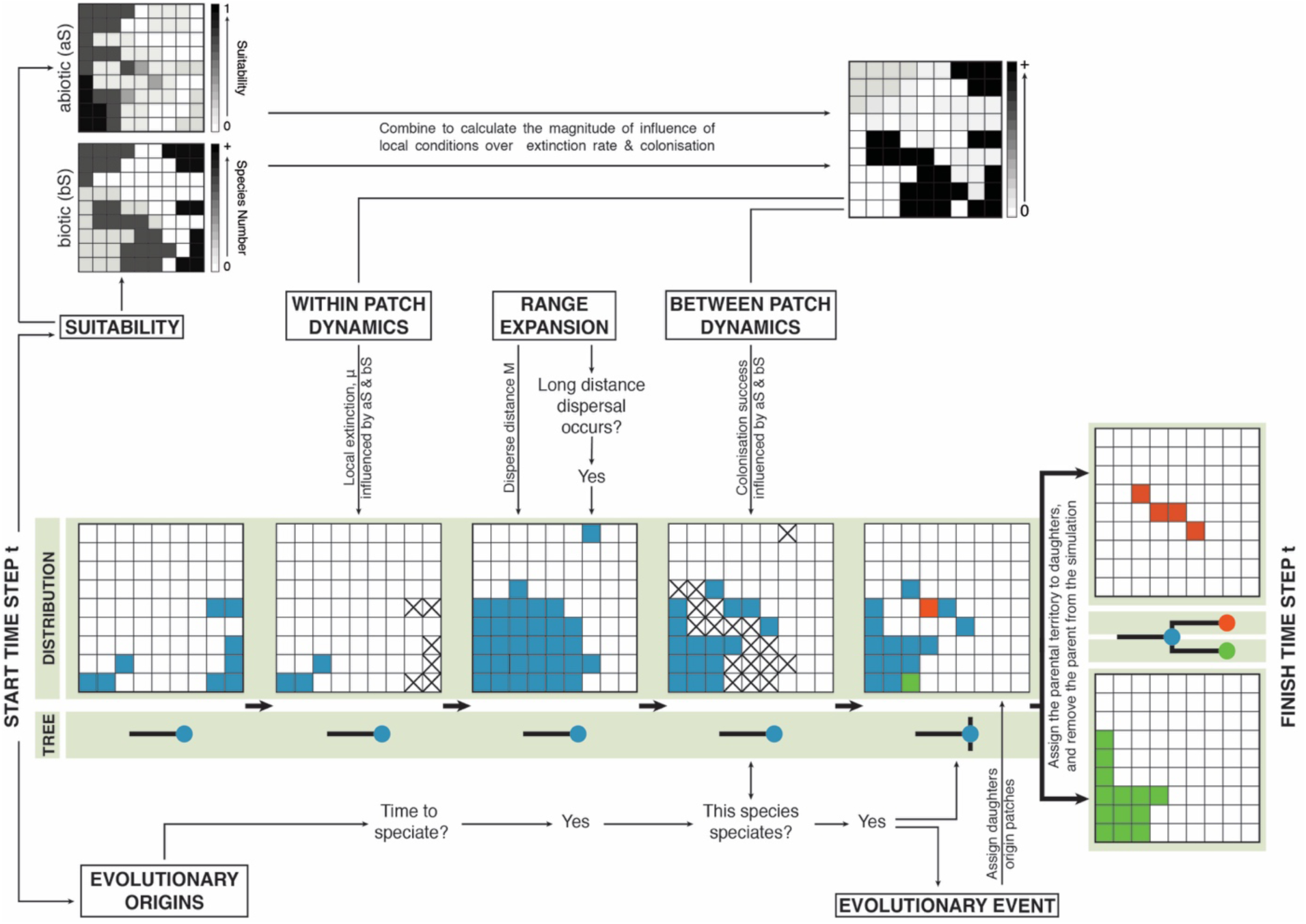
One-time step of the simulation model. Titles bounded by rectangles indicate either the points or the sides of the model overview (the triangle in Figure 1). The grids located on the green band show the territory of a parental species (in blue) and its daughters (red and green). The trees record their evolutionary relationship and time of speciation. For clarity “×” indicates the occurrence of local extinction when applying the influence of “Within Patch Dynamics” and “Between Patch Dynamics” to the territory grid. Here, unsuccessful long-distance dispersal and a speciation event occurred. All the parental territory was successfully divided up between the two daughter lineages.

### Habitat suitability

We considered the landscape suitability of a cell to be combination of two elements or values: a. abiotic suitability, which is dependent on the environmental suitability of the cell for the moving species and on the presence of temporary, slow moving barriers; b. biotic suitability, which is dependent on the number of species present within a cell (Figure 3).

**Figure 3.**
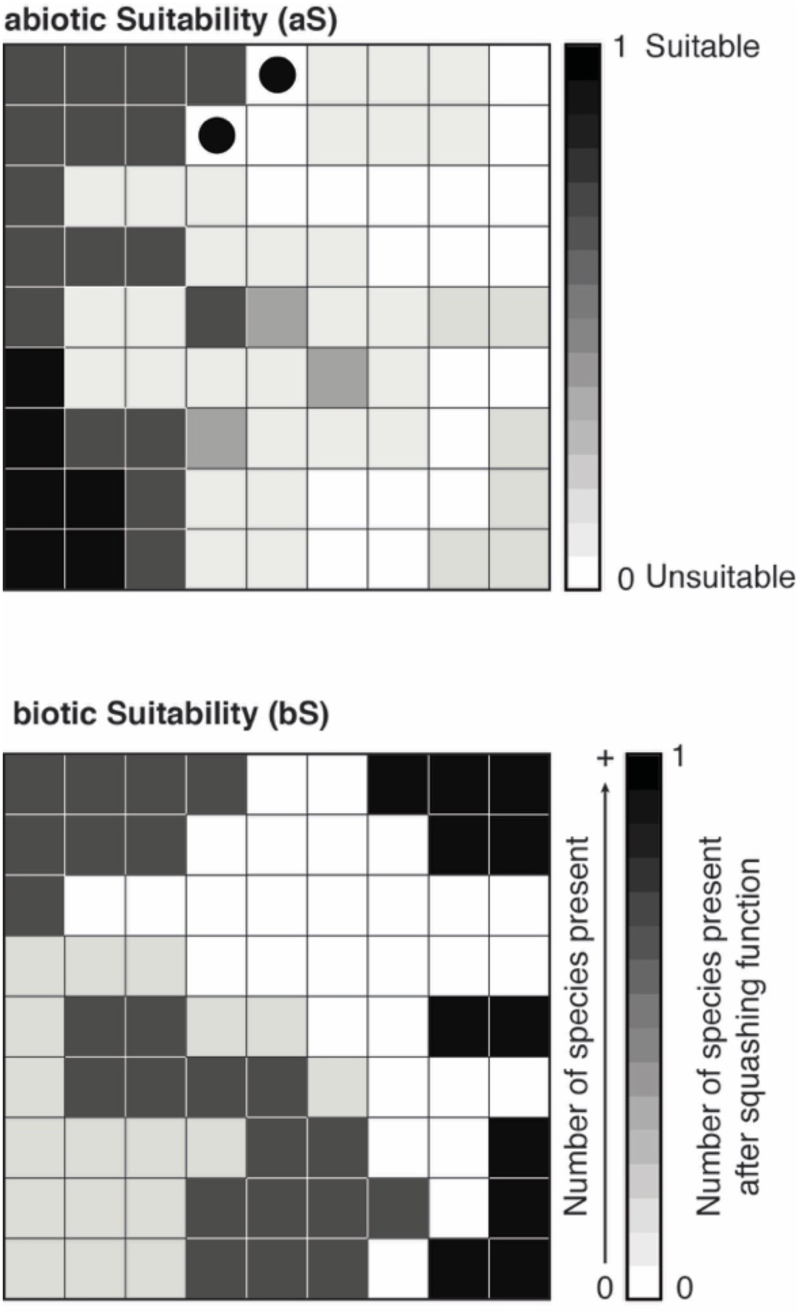
Abiotic and biotic suitability of each cell within the landscape. Suitability is composed of two components: *Abiotic suitability (aS)* probability is created by the environmental suitability (the climatic properties of the cell in relation to the tolerances of the species) and by physical formations (slow moving barriers). *Biotic Suitability (bS)* is the number of species present within a cell (density dependence effects). Abiotic and biotic suitability are bounded between [0,1] values. Environmental abiotic suitability is calculated by an environmental niche model (ENM) that is hindcasted across the time steps *t*. Physical formations are slow moving barriers (mountains, rivers and deserts) that are that appear and disappear over time, and can move to a new geographical space (e.g. mountain building) Physical formations can be represented as cells of low suitability regardless as to whether the cell has high environmental suitability (black circle represents high environmental suitability, however due to a physical formation, the cell is considered to have low suitability).; for simplicity, we consider the landscape to be flat and so barriers are only barriers by width and not height. See more details in the text.

*Abiotic suitability (aS).* This is a suitability probability generated by an environmental niche model (ENM) that has been hindcasted across the time steps of paleoclimate layers. There are a wide variety of models and distances that can be used in ENM hindcasting, including the Mahalanobis distance (Clark et al. 1998), the maximum entropy algorithm (Maxent; Phillips et al. 2006), or Hengl et al. (2009)’s framework combining distances defined by point-pattern analysis and environmental niche factor analysis (ENFA) with a regression-kriging GLM. Each of these models has their advantages and disadvantages (see Chefaoui and Lobo 2008). We use here an approach suggested by Culshaw et al. (2021b), where Hengl et al. (2009)’s method is used to generate an ENM hindcasted projection for each month of the year, and these monthly projections are then overlaid to construct a “yearly” projection. Culshaw et al. (2021b) demonstrated that this approach is more efficient in capturing the general pattern than the usual approach of averaging the results across a collection of months (e.g., Mairal et al. 2017), as the accumulation of errors in the latter can result in the pattern being completely wiped out.

Slow moving barriers are considered to be physical formations, such as mountains, rivers and deserts, that appear and disappear over time, and, depending on the nature of the barrier, can move to a new geographical space, i.e., mountain building and erosion, or river capture, changes the position of a river basin. These physical formations can have a strong influence on the geographical distributions of a species, acting as barriers to or corridors for dispersal and migration, and, hence, deterring or facilitating movement between cells. These barriers can also be included in the *aS* matrix (i.e. values of *aS* per cell in the gridded domain) as low suitability areas. For simplicity, we consider the landscape to be flat, so that barriers are only composed of a width dimension and not of a height dimension. Moreover, we assume that the barrier can be overcome if the species is able to disperse past it to the next suitable cell in one-time step *t*, regardless of whether the terrain cells between a species and the colonised cell are highly unsuitable, hence, trivialising the role played by extreme ecological limitations (e.g. sharks are unable to walk on land and there is only land between them and another cell of water), see Connolly (2005) for the opposite model).

Once the model for estimating abiotic habitat suitability landscape for *t* = 0 for a specific area has been created, we interpolate this model back through time across all the paleoclimate data for the same area. The abiotic landscape suitability’s values (*aS*) lie within the interval [0, 1]. However, as the *aS* matrix can be affected by loss of information caused by events such as population bottlenecking and mass extinction events (Culshaw et al. in 2021a, 2021b) a Brownian motion distribution (Wiener process) is added or subtracted from each *a*S at each time step *t*.

Biotic Suitability (*bS*) refers to the density of the patch in regard to to the number of species. The *bS* matrix contains the number of species within each of the cells. These values are bound within the interval of (0, 1] through the use of a squashing function 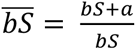, where *a* ∈ ℝ^>0^. Information on the number of species within each cell at any given time is relevant to know the level of interspecific competition present. This knowledge can be used to explain the absence of a given species from regions that are within their set of tolerable environmental conditions (Wiens and Donohue 2004).

### Range evolution

Range evolution describes how individuals disperse through land (Figure 4). If the time step *t* is suitably large (e.g. 500 years), then, in the clear-cut thought process of an invasive species, the land should be completely colonised except for areas that cannot be reached by long distance dispersal, have patch unsuitability, or high competition (Rangel et al. 2007). Here, species in the assemblage share a common spreading dye distribution (Jetz and Rahbek 2001; Colwell et al. 2009; Gotelli et al. 2009), where an individual can move *M* distance in one-time step *t* but can also have long-distance unlimited dispersal to any patch more than *M* distance away with the probability distribution of this event occurring drawn from the Poisson distribution *FD* = *pois*(*λ*_*FD*_) *n*_*FD*_, where magnitude *n*_*FD*_ ∈ ℝ^>0^. Freak dispersal is equally likely of originating in any previously occupied cell.

**Figure 4.**
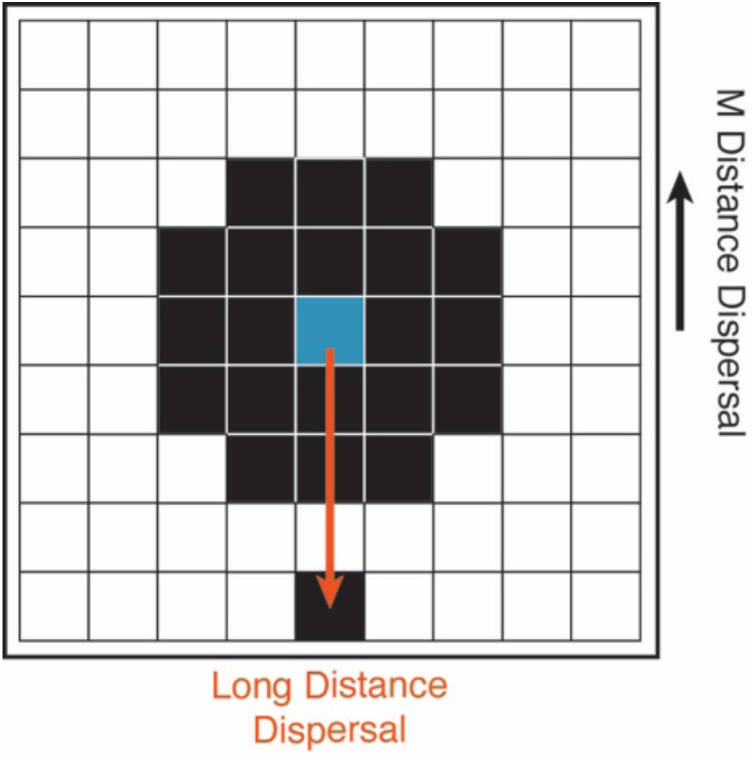
Range Evolution. Here individuals can disperse through the landscape by *M* distance in one-time step *t* or via a long distance and unlimited dispersal to any patch that is greater than *M* distance away. Long distance distribution is equally as likely of originating in any previously occupied cell.

### Evolutionary Origins

During a simulation, this model allows for an evolutionary tree to be built in four different ways (Figure 5). In each of these tree types, the speciation rate (*λ*) and extinction rate (*μ*) must be supplied and the tree is built over a length of time that is given by the length of the simulation (*T*). Parameters *λ* and *μ* can vary over time, i.e., undergo rate shifts, but at any given time *t* rate homogeneity is assumed across lineages, even though this has been shown to be empirically unsound (Rabosky 2014). Since the model assumes phylogenetic niche conservation (i.e., the transmission from ancestor to descendants of the biological and physiological characteristics that determine the fundamental ecological niche of a species (Hutchinson 1957), the probability of a speciation event occurring at time step *t* is only determined by the speciation rate *λ*; this follows a Poisson distribution *pois*(*λ*_)_)*n*_)_, where *λ*_)_ ∈ ℝ^>0^ and magnitude *n*_)_ ∈ ℝ^>0^. The species that undergoes speciation is selected after all species in the simulation have dispersed for that time step t. Once a species has speciated, it is removed from the simulation. The rate of extinction, μ, is drawn from a Poisson distribution *pois*(*λ*_*μ*_)*n*_*μ*_, where *λ*_*μ*_ ∈ ℝ^>0^ and magnitude *n*_*μ*_ ∈ ℝ^>0^. In our model, the Poisson distribution of μ is reshaped by the local patch dynamics occupied by the species (within patch dynamics), which determines the local extinction rate. Ultimately, the accumulation of local extinctions in different patches results in a species global extinction. In addition to these parameters, trees type 2, 3 and 4 require supplying additional information: the speciation time intervals in the case of tree type 2, information on the evolutionary relationships (i.e., the tree topology) for tree type 3; and both the speciation time intervals and information on the phylogenetic relationships for tree type 4. This information adds further constraints upon the building of the tree.

**Figure 5.**
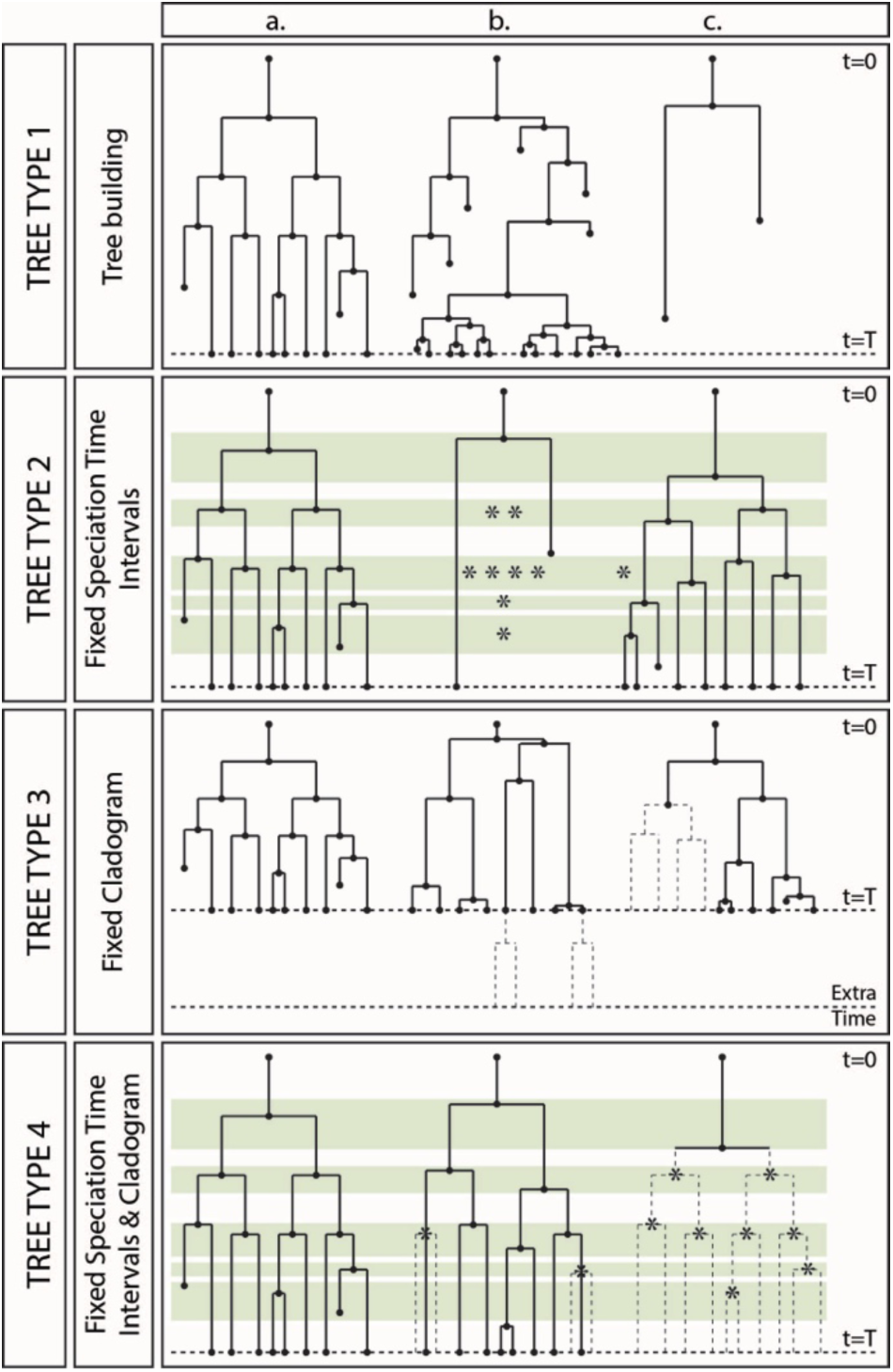
Evolutionary Origins. Evolutionary trees can be built in four different ways, which also represent constraints in the time and position of the speciation event. The figure presents three example trees that have been built under each of these four types for a maximum of *t* = *T* time steps. *Tree type 1* allows simulations to produce any topology and any number of leaves or terminals. There is no cap on species number and there is no constraint in evolutionary relationships, as each species is as equally likely to speciate when it is time to speciate. *Tree type 2* constrains the time interval in which speciation may occur (here presented as a green band), but does not constrain the evolutionary relationships or tree topology, i.e. each species is equally likely to speciate at a given time. Each speciation time interval given covers at least one speciation opportunity (each marked with ∗). For tree “b”, only one speciation event has occurred, with only one of the lineages reaching the present day. Trees “a” and “c” have a similar number of speciation events and the same number of extant, but differ in their tree topology. *Tree type 3* constrains the evolutionary relationships in the tree, i.e., the tree topology. This means that speciation is not constrained to occur within predefined time intervals, but when there is a speciation Figure 5. cont. event, the species that undergoes speciation is predefined. In tree “b”, due to lack of a successful speciation event, one daughter lineage has failed to manifest before the end of the simulation (the missing lineage is denoted by the dotted line); if the tree continues evolving past the simulation time, it may have a tree topology matching that of tree a (*t* = *T* + extra time). In tree “c”, daughters and granddaughters of a species have failed to speciate before the grandparent species became extinct (missing lineage shown by dotted line), and hence tree c can never fully fulfil its predefined topology. *Tree type 4* has the combined constraints of tree types 1 to 3: rates μ and λ, the time interval in which speciation is allowed to take place, and the tree topology. Unlike tree type “3b”, tree type “4b” cannot be given extra time to follow its predefined tree topology (dotted line), as there are no more speciation time intervals during which to speciate. A more extended explanation is given in the text.

Tree type 1 allows simulations to produce any topology and any number of leaves or terminals. There is no cap on species number, and there are no constraints on evolutionary relationships, as each species is equally likely to speciate when it is time to speciate. This tree building method follows the non-equilibrium model assumed by standard birth-death dynamics (Morlon 2014), defined only by *μ* and *λ*. It resembles the method for building evolutionary trees in GSMs suggested by Gotelli et al. (2009).

Tree type 2 constrains the time interval in which speciation may occur (represented in Figure 5 by a green band), but does not set any constraints on the tree topology, i.e. each species is equally likely to speciate when it is time to speciate. This tree building method is similar to the birth-death models that are conditional on a given initiation time (root age) and absolute speciation times, but where the number of terminal species can vary across reconstructed trees (Stadler 2011). Each time interval covers at least one speciation event (marked with ∗). In Figure 5, for example, two of the speciation time intervals contain several possible speciation events. This could be the case when speciation time intervals overlap or are so close that they can be considered to be the same event. It is at the user’s discretion to decide whether single or multiple speciation events are allowed within a single time interval. A speciation event is not required to occur, only that if it were to occur, it can only occur within the speciation time intervals. Therefore, the absolute value for time to speciation (branch length) is only partially predefined. For tree “b” only one speciation event has occurred, with only one of the descendant lineages reaching the present day. This implies that speciation failed to occur for each of the daughter lineages within a speciation time interval before either the descendant became extinct or the simulation ended. Trees “a” and “c” have a similar number of speciation events (9 and 8 respectively) and the same number of extant species in the tree; however, they differ in their topologies (Figure 5).

Tree type 3 constrains the evolutionary relationships in the tree, i.e., the tree topology. This means that speciation is not constrained to occur within predefined time intervals, but when there is a speciation event, the species that undergoes speciation is predefined. As in tree type 2, a species is not dictated to speciate; hence, the absolute value for time to speciation (branch length) is not considered. However, should a species fail to speciate, then all its possible daughters will not speciate either, i.e. a descendant speciation event cannot precede that of its ancestor. In Figure 5, tree “b”, due to lack of a successful speciation event, one daughter lineage has failed to manifest before the end of the simulation (the missing lineage is denoted by a dotted line). In theory, should tree “b” be given extra simulation time, it may exhibit a tree topology matching that of tree a. In tree “c”, daughters and granddaughter lineages of one of the initial species have failed to speciate before the grandparent species became extinct (missing lineage shown by dotted line), and hence, tree “c” can never fully fulfil its predefined tree topology (Figure 5).

Tree type 4 has the combined constraints of tree types 1 to 3: rates *μ* and *λ*, the time interval in which speciation is allowed to take place, and the tree topology. Thus, as in tree type 2, the absolute value for time to speciation (branch length) is predefined. Unlike tree type “3b”, tree type “4b” cannot be given extra time to follow its predefined tree topology (dotted line), as there are no more speciation time intervals during which to speciate (Figure 5).

Having these different tree types is advantageous. If the user has more information than just *λ* and *μ*, he can make a more informative simulation including lineage divergence times or tree topology, and this allows for a more direct comparison with empirical or previously simulated phylogenies. On the other hand, if the user has doubts or lacks estimations of *μ* and/or *λ*, then, information about only the topology can be used to explore the *μ* and *λ* parameter space to find a combination of these two rates that works. This is particularly interesting because estimations of *μ* are in general considered to be problematic to obtain (Sanmartin and Meseguer 2016), while estimates of *λ* and speciation times are deemed reliable.

### Extinction

Extinction (local, eventually resulting in global) and successful colonisation can be important explanations for the absence of a clade from a given area. Although extinction needs not directly involve dispersal, the absence of a clade from a region owing to extinction or unsuccessful (re)colonisation indicates a limitation on dispersal (Wiens and Donohue 2004). In our model, extinction and colonisation success following dispersal are not equiprobable among grid cells; instead, they are dependent on the particular values the environmental variables adopt in each cell.

### Within and between patch dynamics

Within and between patch dynamics models the relationship between habitat suitability and extinction rate (*μ*), and suitability and the probability of colonisation success (ℙ(*C*)) (Figure 6). ℙ(*C*) may undergo rate shifts through the simulation. It is known that the probability of extinction is dependent on abiotic or biotic factors (i.e., passive replacement and active displacement, Silvestro et al. 2015). This refers to the local extinction of individuals within species; if all individuals of a species are removed, then a global extinction occurs and the species can no longer speciate and leave behind daughter lineages. In our model, we combine the abiotic suitability matrix (*aS*) with the biotic suitability matrix (*bS*) (Figure 6a) in order to calculate the magnitude of the influence of the local environmental conditions in the patch (*i*, *j*) at time step *t* over the Poisson distribution *μ* and ℙ(*C*) are drawn from, separately; this allows calculation of the local extinction rate and colonisation success (Figure 6b).

**Figure 6.**
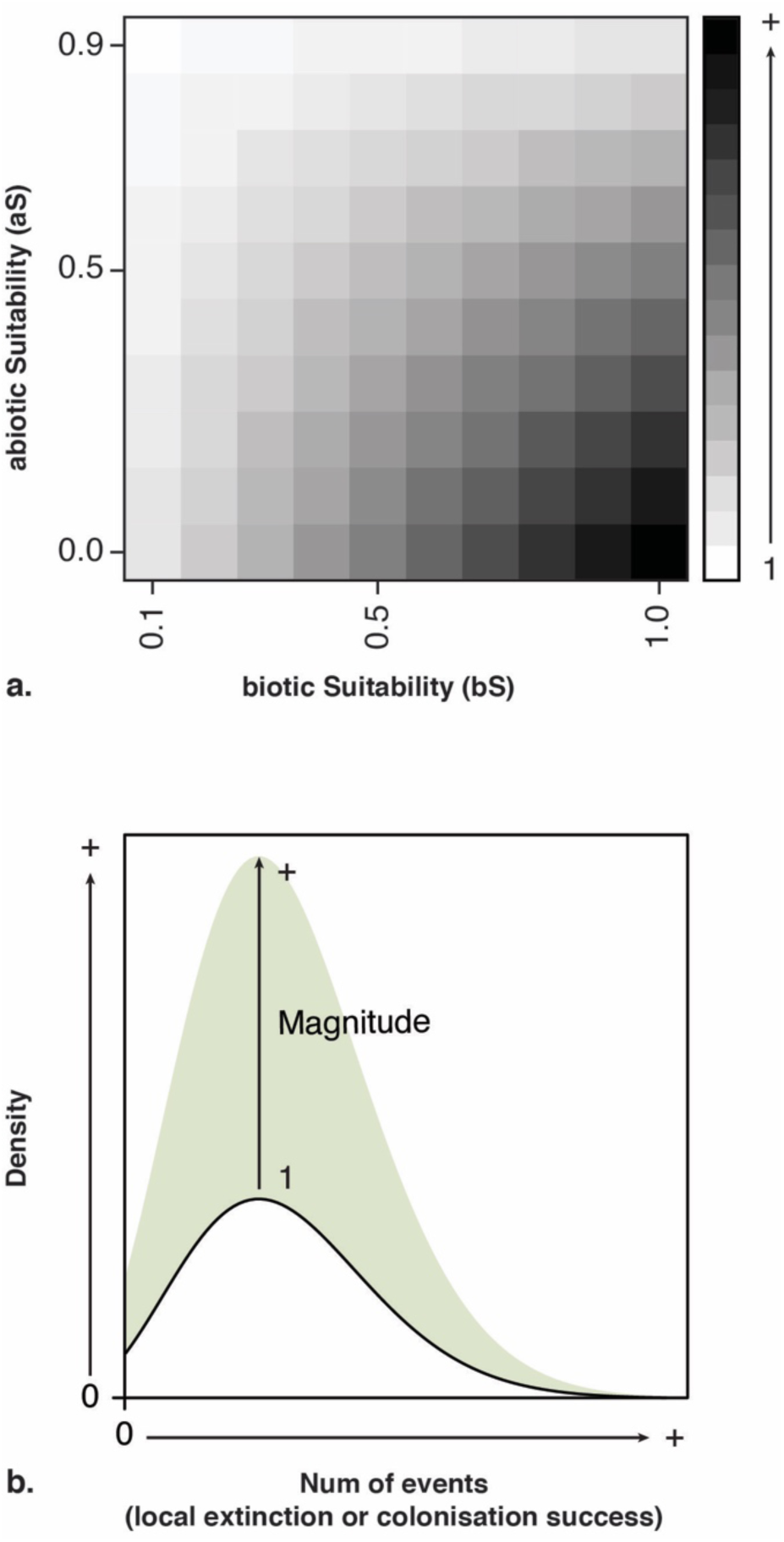
Demonstrating how local background extinction rate is calculated from values of cell values of abiotic and biotic values at time. ***t***. a. The model default to combine *aS* and 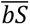 together is linear (1 − *aS*)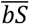, where 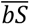 = shaper function(*bS*) member (0, 1] as there is always at least one species present in a patch that local background extinction is calculated for and *aS* ≠ 1 as conditions can never be 100% perfect. The default magnitude is exponential and its gradient is controlled by *n* ∈ ℝ^>0^. As *n* → 0, the influence of the magnitude over the *μ* Poisson distribution becomes minimal to non existent, e.g. each patch has enough resources for all. a. shows the default magnitude with any *n*. Notice that as *aS* → 1 and *bS* → 0 then the magnitude → 1 i.e. the *μ* Poisson distribution and colonisation success remain unchanged.

The default model to combine *aS* and 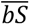 together is linear (1 − *aS*)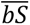, where 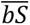 = shaper function(*bS*) member (0, 1], as there is always at least one species present in a patch for which local extinction is calculated, and *aS* ≠ 1 as conditions can never be 100% perfect. The default magnitude is exponential and its gradient is controlled by *n* ∈ ℝ^>0^. As *n* → 0, the influence of the magnitude over the *μ* Poisson distribution becomes minimal to non-existent, e.g. each patch has enough resources for all species. Figure 6a shows the default magnitude with any *n*. As *aS* → 1 and *bS* → 0, the magnitude → 1 i.e. the *μ* Poisson distribution and colonisation success remain unchanged. Figure 6b presents the Poisson distribution for local extinction μ at patch (*i*, *j*) at time *t*:

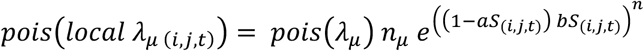

The probability for local colonisation success at patch (*i*, *j*) at time *t* is:

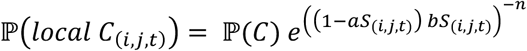

The user can change all default models.

### Division of parental territory

An Evolutionary Event occurs when Lineage Information tells the model that a speciation event has occurred, once that species has dispersed (Figure 2). In our model, the Evolutionary Event component of the model addresses the geographical mode of speciation by assigning origin patches, chosen from the parent’s territory, to the daughters (in Figure 7, these are denoted in black, and with red and green dots, respectively). All patches that form the parental territory are equally likely to be selected. After this, the parent’s territory is divided between the two daughter lineages. There are three ways a parental territory can be divided into its daughters. Notice that only territory that is *M* distance away from the origin patch is considered; any unreachable territory is disregarded, i.e. black patches lacking a green or red dot in Figure 7.

a. Daughters originate from the same patch; both daughters will inherit the same territory.
b. Daughters originate from different patches and do not have the possibility to inherit the same territory. The patches lie M distance away from only one origin patch. The territory is divided up without conflict.
c. Daughters originate from different patches but have the possibility to inherit the same territory. In this case, the territory will be divided up, with patches M distance away from both origin patches being assigned to the daughter that is closer. If the daughters are the same distance away from the patch being assigned, the conflict is resolved by pseudo randomly assigning it to a daughter.

**Figure 7.**
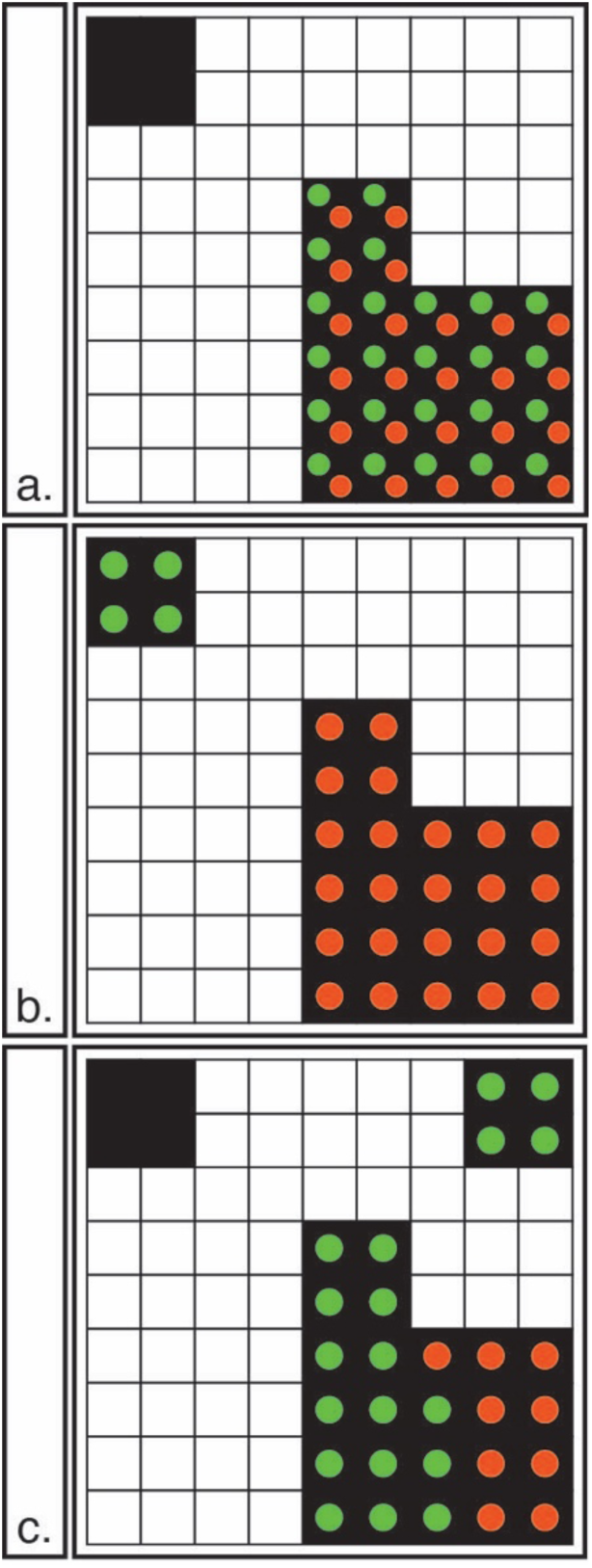
Range inheritance scenarios: The figure shows the division of the parental territory between the daughters at the time of an evolutionary event. The parent’s territory is denoted in black, with each daughter denoted as a red or green dot. There are three ways in which parental territory can be assigned to the daughters. a. Daughters originate from the same cell. Both daughters will inherit the same territory. b. The cells lie M distance away from only one origin patch. The territory is divided up without conflict. c. Daughters originate from different cells but have the possibility to inherit the same territory. In this case the territory will be divided up, with patches *M* distance away from both origin cells being assigned to the daughter that is closer. If the daughters are the same distance away from the cell being assigned, the conflict is resolved by pseudo randomly assigning it to a daughter.

Once a parent species has speciated, it is removed from the simulation. The three models of fragmentation of the parental territory represent the sympatric, parapatric or peratric/allopatric speciation modes (Gotelli et al. 2009).

## Conclusions and Some Suggestions for Testing the Model

Rangel et al. (2007) evaluated the robustness of their GSM model in terms of its ability to predict long-discussed tenets in biogeography, such as the latitudinal diversity gradient or the mid-domain effect based on the distribution of 3000 species of South American birds. Predicted spatial patterns closely resemble observed ones and proved sensitive to niche dynamics processes. This allowed them to evaluate the roles of both evolutionary and ecological processes in explaining spatial patterns in species richness and geographic range size distribution, thus providing a “link between ecology and historical biogeography under integrated theoretical and methodological frameworks”. Their GSM model was novel in using (artificially) fluctuating environmental conditions and a single ancestral origin for the species generated in the model, thus providing a link between speciation events.

In our general simulation model, we provide a link between speciation events in different way by instead of focusing on species richness patterns or the predicted spatial distribution of species (a pattern-oriented modelling approach (Grimm et al. 1996, 2005), we propose to test our model by explicitly introducing the evolutionary component in the form of a phylogeny and associated lineage divergence times. We select parameter values for the final simulation based on maximising the similarity between observed and simulated spatial patterns, specifically: i. the geographic distribution of extant species; and ii. the resultant phylogenetic pattern from the simulations. Accounting for phylogenetic relationships and a time dimension in comparing the simulated and the real world is actually important to increase the realism of the model and its power to detect system-level properties (Gotelli et al. 2009). The discipline of biogeography has both spatial and temporal dimensions, and the level of niche conservatism can change over the history of a lineage (Sanmartín 2014). The shape and length of a phylogeny can also be informative about the actual processes involved in their generation in the first place: for example, “broom- and-handle shape” phylogenies are typically associated with high background extinction rates or mass extinction events (Sanmartín and Meseguer 2016). Hence, introducing phylogenetic shape and time in the simulation outcomes might help us test the role played by background extinction and high extinction rates in the formation of both spatial patterns and evolutionary patterns.

Other differences between our model and Rangel et al. (2007) lies in the modelling of extinction and the introduction of actual empirical data in the model. In their GSM model, speciation is caused by fragmentation and isolation, while extinction is linked to reductions in range size up to a certain threshold. In our model, speciation is brought about via a speciation rate, and extinction by an extinction rate that is dependent on abiotic (palaeoenvironmental (climate)) and biotic (density-dependence) factors, thus helping introduce a basic species competition to the model. Also in our GSM, the speciation event can be further informed with speciation time intervals and the empirical tree cladogram. This allows for a more direct comparison of the GSM results to an empirical example, but also interestingly, this lends to an opportunity to explore the speciation and extinction rate space to find suitable pairings of speciation and extinction rates for phylogenies that are unsuitable for traditional birth-death model. Rather than using a cyclical environmentally fluctuating landscape as in Rangel and collaborators, we use a time series of palaeoclimatic layers spanning the last 20 million years. Using empirical data, our understanding of the evolutionary relationships and biogeographic history of real organisms (Gotelli et al. 2009) can help better refine and constrain the range of parameter values examined during the simulations. This might be especially important when we do not have a large phylogenetic and distribution dataset as input.

One nice example of such a case comes from a biogeographic pattern of plant distributions in Africa tackled by our team. The Rand Flora (RF) pattern is a continental-scale geographic disjunction in which sister species are distributed on opposite sides of the African continent: in Macaronesia-northwestern Africa, Eastern Africa-southern Arabia and southern Africa (Sanmartín et al. 2010). Dated phylogenies and biogeographic analyses in some of these groups suggest that most RF disjunctions date from the Early Miocene to the Mid Pliocene periods, concurrent with an aridification trend in Africa, a result of plate tectonics and mountain building (Pokorny et al. 2015). Species populations are often small in size and have highly restricted distributions. Recent ENM reconstructions (Mairal et al. 2017b) suggest that RF-lineage distributions were broader in the past and that range expansion was achieved by climatic corridors, which were later interrupted by successive aridity waves. One difficulty with inferring the origin of the RF biogeographic pattern is that most RF phylogenies comprise of less than 5-10 species and often exhibit “broom-and-handle” shapes, with long stem-branches and young crown clades, indicative of high extinction rates (Sanmartín and Meseguer 2016). The long-time intervals without cladogenetic events along the phylogeny hinders the use of birth-death macroevolutionary and biogeographic inferential models. Culshaw et al. (2021a, 2021b) handled this uncertainty by adopting an integrative inferential approach, combining information from biogeographic analyses and ENMs to disentangle the processes behind the RF pattern in the small (only three species) but highly disjuct (continental distribution) genus *Camptoloma* (Scrophulariaceae). As in many RF lineages, information on species and population distributions is scarce.

The spatially-explicit forward time computational simulation model could be also another avenue to tackle the RF pattern. For example, one could use our GSM model and paleoclimate data from Africa (as done in Culshaw et al. 2021a, 2021b, and Mairal et al. (2017) to try and recreate the Rand Flora distribution pattern (continental disjunction between the margins of Africa) and the evolutionary phylogenetic shapes characterising RF lineages (ancient stem-age, young crown-group and long internodes). One advantage of using the GSM approach is that we can let estimates of diversification and background extinction rates obtained from the birth-death framework (Pokorny et al. 2015) to inform or constrain the range of values adopted by speciation and extinction rates in the model), thus increasing the realism of the model in relation to the examined pattern, but also helping to reduce the uncertainty in the comparison between observed and simulated patterns. One strong assumption of our model is that we assume niche conservatism over the entire simulation. Species expand their range occupying adjacent cells with similar conditions. Rangel et al. (2007)’s GSM allowed niche evolution, by modelling random small deviations, from the niche breadth centre using a Brownian model (sigma). Assuming niche conservatism is risky for long geological time scales. For the Rand Flora, this is not such a crucial impediment because biogeographic inferences seem to point out to a conservation of ancestral tolerances and fragmentation being responsible for the RF distribution (Rincón-Barrado et al. 2021; Culshaw et al. 2021a, 2021b.); it is also a more conservative approach in the absence of information. Moreover, Rangel et al. (2007)’s model, initial tolerance to climatic conditions is determined by the initial location of the first simulated species (i.e., climatic conditions of the starting cell). In our model, this would be given by the actual ENM models based on empirical data and by the real palaeoclimate layers for our specific biogeographic domain (Africa), thus again reducing uncertainty in the model.

In sum, we argue that the computational general simulation model presented here could be an interesting alternative to explore hypotheses on the spatial patterns and processes underlying general distribution biodiversity patterns, especially when the initial input data is small, thereby introduces large uncertainties in the model.

## Declaration

This work was created as the 3^rd^ chapter of the Ph.D. thesis of the leading author V.C., entitled “Developing new tools to address the impact of climate change on the evolutionary and distributional history on plant lineages”, successfully defended on 28^th^ February 2020.

## Acknowledgments

We thank Andrew Hendry for the helping with the initial concept of the model, Dr Joaquín Hortal for providing the introduction to Thiago Rangel. Thiago Rangel for welcoming V.C. into LETS, Universidade Federal de Goiás, Brasil. V.C. was funded by Ministerio de Economía y Competitividad MINECO Ph.D. (FPI) Fellowship BES-2013-065389, supervised by I.S.. I.S. was funded by the Spanish Government and European Regional Development Fund, grant CGL2015-67849-P (MINECO/FEDER).

## Author contributions

VC designed the model under the guidance of TFLVBR and help from IS. VC and IS co-wrote the manuscript.

## Author Approvals

The authors declare that all authors have seen and approved the manuscript, and that it hasn’t been accepted or published elsewhere.

## Conflict of Interest

The authors declare that the research was conducted in the absence of any commercial or financial relationships that could be construed as a potential conflict of interest.

## Notes

### Competing Interest Statement

The authors have declared no competing interest.

### Summary of Updates

Within the manuscript, Isabel Sanmartin's affiliation was incorrect. It has now been updated

